# CoxMDS: Multiple Data Splitting for High-dimensional Mediation Analysis with Survival Outcomes in Epigenome-wide Studies

**DOI:** 10.1101/2025.10.11.681836

**Authors:** Minhao Yao, Peixin Tian, Xihao Li, Shijia Bian, Gao Wang, Yian Gu, Ana Navas-Acien, Badri N. Vardarajan, Daniel W. Belsky, Gary W. Miller, Andrea A. Baccarelli, Zhonghua Liu, the Alzheimer’s Disease Neuroimaging Initiative

**Affiliations:** Department of Statistics and Actuarial Science, The University of Hong Kong, Pokfulam, Hong Kong SAR, China; First Affiliated Hospital of Kunming Medical University, Yunnan, China; Department of Biostatistics, University of North Carolina at Chapel Hill, Chapel Hill, NC, USA; Department of Genetics, University of North Carolina at Chapel Hill, Chapel Hill, NC, USA; Department of Biostatistics and Bioinformatics, Emory University, Atlanta, GA, USA; Center for Statistical Genetics, Gertrude H. Sergievsky Center, Columbia University Medical Center, New York, NY, USA; Department of Neurology, Columbia University, New York, NY, USA; Taub Institute on Alzheimer’s Disease and the Aging Brain, Columbia University, New York, NY, USA; Department of Epidemiology, Joseph P. Mailman School of Public Health, Columbia University, New York, USA; Department of Environmental Health Sciences, Columbia University, New York, NY, USA; Butler Columbia Aging Center, Columbia University Mailman School of Public Health, New York, USA; Office of the Dean, Harvard T.H. Chan School of Public Health, Boston, MA, USA; Department of Biostatistics, Columbia University, New York, NY, USA

**Keywords:** Omics mediation analysis, False discovery rate, Survival outcomes, DNA methylation, Data splitting

## Abstract

Causal mediation analysis investigates whether the effect of an exposure on an outcome operates through intermediate variables known as mediators. Although progress has been made in high-dimensional mediation analysis, current methods do not reliably control the false discovery rate (FDR) in finite samples, especially when mediators are moderately to highly correlated or follow non-Gaussian distributions. These challenges frequently arise in DNA methylation studies. We introduce CoxMDS, a multiple data splitting method that uses Cox proportional hazards models to identify putative causal mediators for survival outcomes. CoxMDS ensures finite-sample FDR control even in the presence of correlated or non-Gaussian mediators. Through simulations, CoxMDS is shown to maintain FDR control and achieve higher statistical power compared with existing approaches. In applications to DNA methylation data with survival outcomes, CoxMDS identified eight CpG sites in The Cancer Genome Atlas (TCGA) that are consistent with the hypothesis that DNA methylation may mediate the effect of smoking on lung cancer survival, and two CpG sites in the Alzheimer’s Disease Neuroimaging Initiative (ADNI) that are consistent with the hypothesis that DNA methylation may mediate the effect of smoking on time to Alzheimer’s disease conversion.

## 1 Introduction

Causal mediation analysis aims to understand the causal pathway from an exposure to an outcome of interest through an intermediate mediator variable [1–4], and has become a valuable tool in genome-wide epigenetic studies, where DNA methylation CpG sites mediate the relationships between modifiable exposures and health outcomes [5–9].

DNA methylation involves the covalent bonding of a methyl group (CH3) to the C5 position of cytosine, which plays a regulatory role in gene expression by either interacting with relevant genes or obstructing the binding of transcription factors to DNA [10]. Since DNA methylation is a reversible biochemical process [11], targeting DNA methylation may offer novel therapeutic opportunities for disease prevention and intervention [12, 13].

On Illumina Infinium DNA methylation microarrays (such as the HumanMethylation450 and EPIC BeadChips), methylation at each CpG site is quantified using paired probe intensities for methylated (*Methy*) and unmethylated (*Unmethy*) signals. The methylation level is usually summarized as either the *β*-value, *β* = *Methy/*(*Methy* + *Unmethy* + *α*), which ranges from 0 to 1 with a small offset *α*, or the *M*-value, a logit-like transformation of the *β*-value [14–16]. *β*-values are often modeled using beta-family distributions [17, 18], whereas *M*-values, even after transformation, are not guaranteed to follow a Gaussian distribution [19]. Furthermore, methylation levels show strong local correlation among neighboring CpGs, a phenomenon known as co-methylation [20].

As a result, array-based DNA methylation data are typically non-Gaussian and highly correlated, which creates major challenges for statistical modeling and limits the effectiveness of existing methods for high-dimensional omics mediation (xMediation) analysis. In particular, it remains difficult to reliably identify CpG sites that mediate the effect of an exposure on a survival outcome while controlling the false discovery rate (FDR) [21] in finite samples. Several approaches have extended the Cox proportional hazards model [22] to accommodate high-dimensional mediators. For example, Luo et al. [7] combined sure independence screening (SIS) [23], the minimax concave penalty (MCP) [24], and the Sobel test [25] to select mediators from DNA methylation data, and Zhang et al. [26] proposed a de-biased Lasso estimator for survival outcomes. However, neither method provides finite-sample FDR control. More recently, Tian et al. [27] introduced the CoxMKF method, which aggregates multiple knockoffs to achieve finite-sample FDR control. Yet, the knockoff filter may perform poorly when mediators are moderately to highly correlated [28], and because knockoff copies are generated under a multivariate Gaussian assumption, power can decline when the mediators deviate from Gaussianity [28]. These limitations highlight the need for new methods that guarantee finite-sample FDR control while retaining power in the presence of correlated and non-Gaussian mediators.

In this paper, we propose CoxMDS, a new method for identifying DNA methylation CpG sites that mediate the effect of an exposure on a survival outcome. Building on the multiple data splitting strategy of Dai et al. [28], CoxMDS guarantees finite-sample FDR control even when mediators are moderately to highly correlated or follow non-Gaussian distributions. The procedure consists of three main steps:

**Step 1. Candidate mediator filtering:** Following prior work [7, 27], we fit a linear regression model for each mediator to test the exposure–mediator association. The resulting *p*-values are adjusted using the Benjamini–Hochberg (BH) procedure [21] to retain the most significant candidate mediators.
**Step 2. Mirror statistic construction:** The data are randomly split into two groups. In Group I, we fit a Cox proportional hazards model with the minimax concave penalty (MCP) to the candidate mediators from Step 1 and retain those with non-zero coefficients. In Group II, we refit the Cox model including only the selected mediators from Group I. For each mediator, a mirror statistic is then computed by combining the coefficients from both groups. This process is repeated multiple times to stabilize the results.
**Step 3. Mediator selection with finite-sample FDR control:** For each candidate mediator, we calculate its inclusion rate [28], which measures its frequency of selection across data splits. A data-adaptive threshold is applied to the inclusion rates to select mediators at the target FDR level.

Through extensive simulations, we show that CoxMDS guarantees finite-sample FDR control and achieves higher power than CoxMKF when mediators are non-Gaussian or moderately to highly correlated. In real data applications, CoxMDS identified eight CpG sites that may mediate the effect of smoking on lung cancer survival in The Cancer Genome Atlas (TCGA) cohort, and two CpG sites that may mediate the effect of smoking on time to Alzheimer’s disease conversion in the Alzheimer’s Disease Neuroimaging Initiative (ADNI) cohort.

## 2 Methodology

### 2.1 High-dimensional mediation model with survival outcomes

Suppose we have an i.i.d. sample of size *n*. For subject *i* (*i* = 1, …, *n*), let *D*_*i*_ denote the true event time, *C*_*i*_ the censoring time, and *T*_*i*_ = min(*D*_*i*_, *C*_*i*_) the observed survival time with event indicator Δ_*i*_ = *I*(*D*_*i*_ ≤ *C*_*i*_). We aim to investigate the causal mechanism from an exposure *X*_*i*_ to the survival outcome *T*_*i*_ through a *p*-dimensional mediator vector ***M***_*i*_ = (*M*_*i*1_, …, *M*_*ip*_)^⊤^, as illustrated in Figure 1.

**Figure 1:**
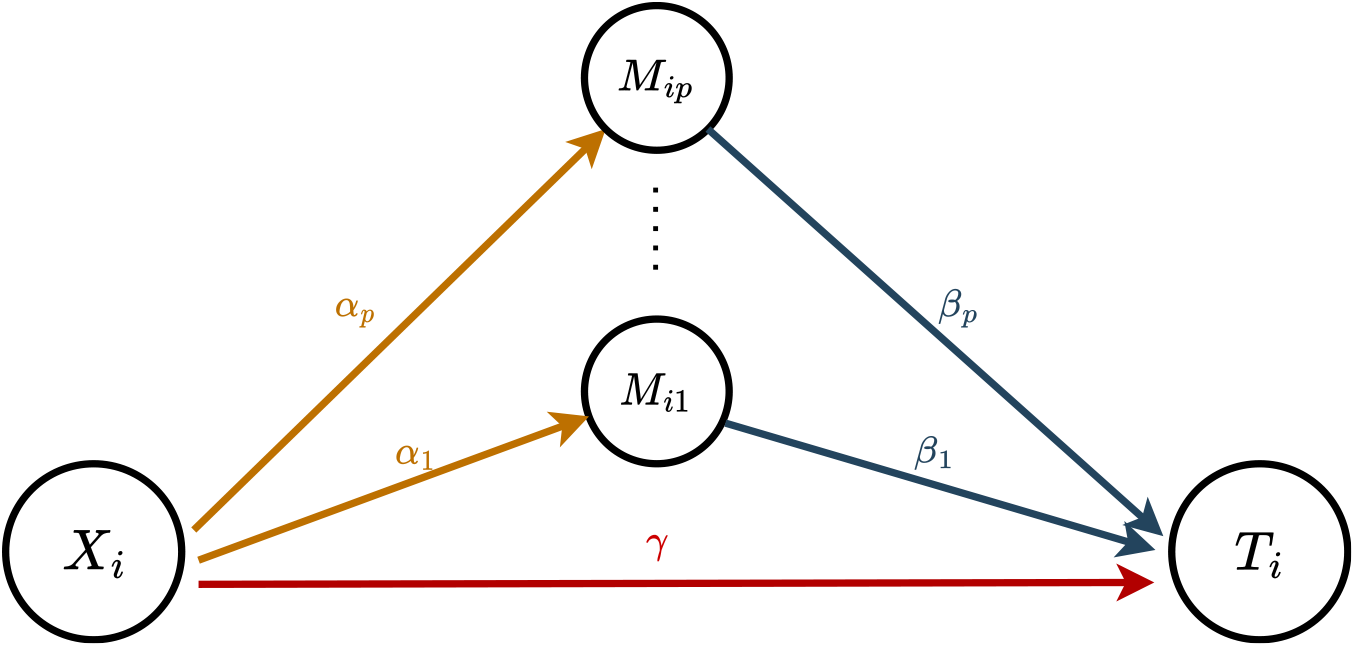
Causal diagram of multiple mediators related with the exposure variable *X*_*i*_ and the survival outcome *T*_*i*_.

Specifically, we consider the following two models for the mediators and survival outcome [3, 4, 27]:

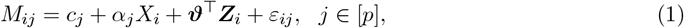

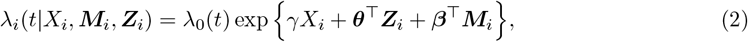

where [*p*] denotes the set of {1, 2, · · ·, *p*}, ***Z***_*i*_ = (*Z*_*i*1_, · · ·, *Z*_*iq*_)^⊤^ is a *q*-dimensional baseline covariate vector (e.g., sex, age), *ε*_*ij*_ is the error term, and *λ*_*i*_(*t*|*X*_*i*_, ***M***_*i*_, ***Z***_*i*_) and *λ*_0_(*t*) are the Cox proportional hazards model and an unspecified baseline hazard function, respectively. In model (1), let ***α*** = (*α*_1_, · · ·, *α*_*p*_)^⊤^ denote the regression parameter vector relating the exposure to the mediators, and let ***c*** = (*c*_1_, · · ·, *c*_*p*_)^⊤^ denote the vector of intercepts. In model (2), *γ* is the direct effect of the exposure on the outcome, and ***β*** = (*β*_1_, · · ·, *β*_*p*_)^⊤^ is the effect of the mediators on the survival outcome adjusting for the effect of the exposure. ***ϑ*** and ***θ*** are the regression coefficients relating the covariates to the mediators and to the survival outcome, respectively. Let *H*_1,*α*_ = {*j* ∈ [*p*] : *α*_*j*_≠ 0} denote the set of mediators with non-zero exposure-mediator associations, and let *H*_1,*β*_ = { *j* ∈ [*p*] : *β*_*j*_ ≠0 } denote the set of mediators with non-zero mediator-outcome associations. Then, *H*_1_ = *H*_1,*α*_ ∩ *H*_1,*β*_ denotes the set of true mediators, and *H*_0_ = {*j* ∈ [*p*] : *j* ∈*/ H*_1_} is the set of non-mediators.

Under the potential outcomes framework [29, 30], we can decompose the effect of the exposure on the survival outcome using the difference in counterfactual log hazards [3, 31]. Let *T* (*x*, ***m***) denote the survival time when the exposure is set to *x* and the vector of mediators is set to ***m, M*** (*x*) = (*M*_1_(*x*), · · ·, *M*_*p*_(*x*))^⊤^ denote the vector of mediators when the exposure is set to *x*, ***Z*** denote the baseline covariates, and *x*^∗^ denote the reference level of the exposure. The total effect (TE) of the exposure on the outcome, the natural direct effect (NDE), and the natural indirect effect (NIE) are defined respectively as follows [3, 31]:

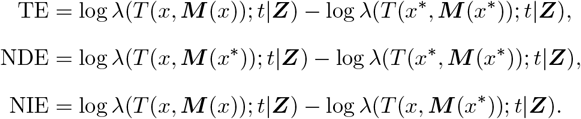

According to the definitions, we can decompose the total effect as TE = NDE + NIE.

As noted in VanderWeele [3], we require the following four standard assumptions for the identification of NDE and NIE in models (1) and (2):

(A1) *T* (*x*, ***m***) ⊥ *X*|***Z***: there is no unmeasured confounding between the exposure and the survival outcome.

(A2) *T* (*x*, ***m***) ⊥ *M*_*j*_|*X*, ***Z*** for *j* ∈ [*p*]: there is no unmeasured confounding between mediators and the survival outcome.

(A3) *X* ⊥ *M*_*j*_|***Z*** for *j* ∈ [*p*]: there is no unmeasured confounding between the exposure and mediators.

(A4) *T* (*x*, ***m***) ⊥ *M*_*j*_(*x*^∗^)|***Z*** for *j* ∈ [*p*]: there is no exposure-induced confounding between the mediators and the survival outcome.

If assumptions (A1) - (A4) hold, then we have the following approximation for the counterfactual log hazard [3, 27, 31]:

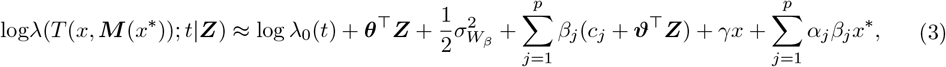

where 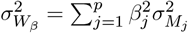, and 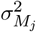 is the variance of the *j*th mediator. Therefore, in models (1) and (2), we have the following expressions of NDE and NIE, respectively:

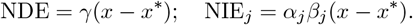

### 2.2 CoxMDS Framework

In high-dimensional mediation analysis, the number of candidate mediators *p* is often much larger than the sample size *n*, so the traditional Cox proportional hazards model (2) cannot be directly applied. Recently, Tian et al. [27] proposed a framework, “CoxMKF,” which integrates the Benjamini–Hochberg (BH) procedure [21] with the aggregation of multiple knockoffs [32] to achieve finite-sample FDR control in high-dimensional mediation analysis with survival outcomes. However, the knockoff filter requires exact knowledge of the joint distribution of mediators. When mediators are moderately to highly correlated or follow non-Gaussian distributions, the knockoff filter may suffer from reduced power in identifying true mediators [28]. To overcome this limitation, we propose “CoxMDS,” a novel framework that adapts the idea of multiple data splitting [28]. CoxMDS accommodates correlated or non-Gaussian mediators while maintaining finite-sample FDR control in high-dimensional mediation analysis with survival outcomes, as illustrated in Figure 2.

**Figure 2:**
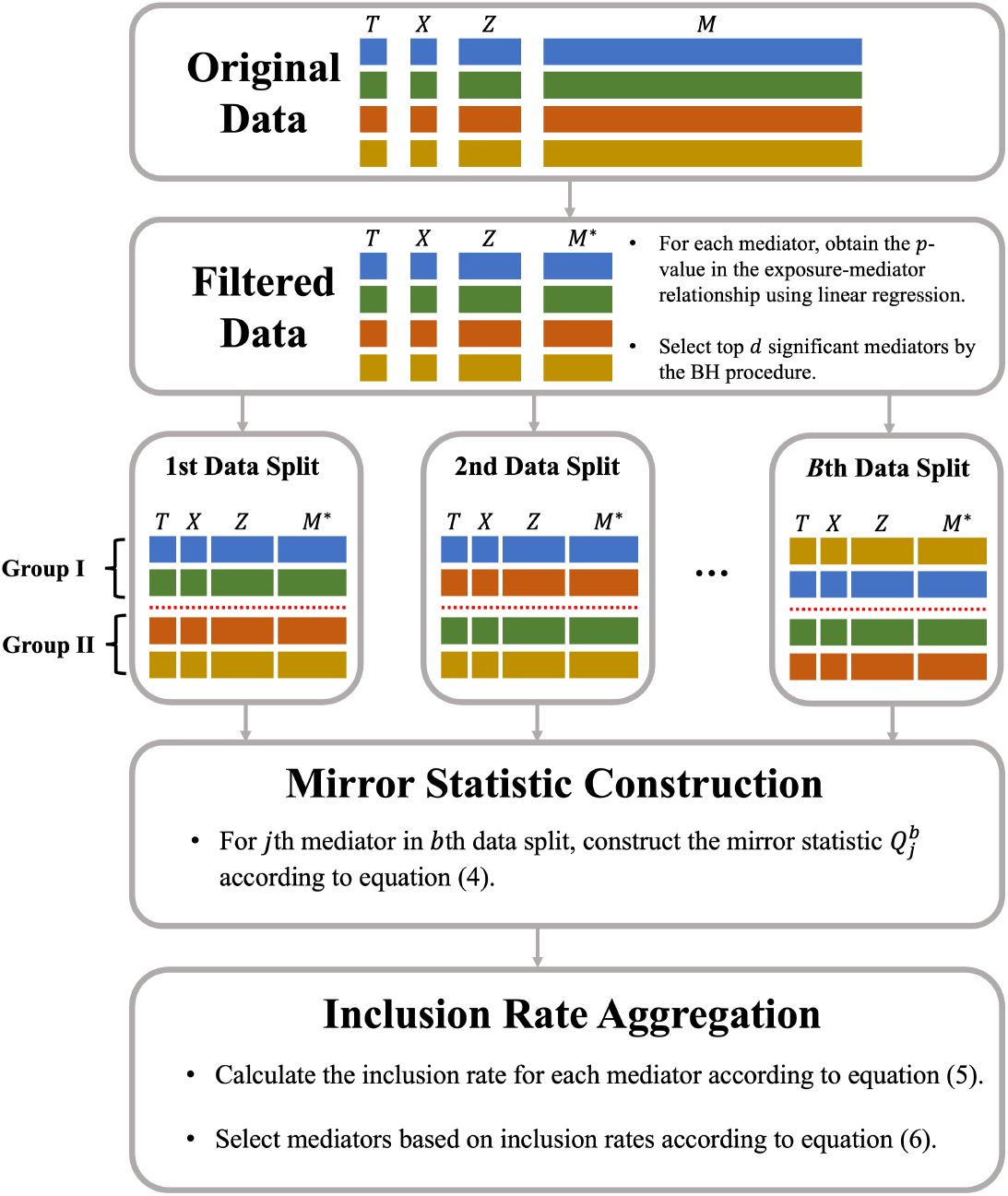
Framework of CoxMDS. First, we apply the BH procedure to the exposure–mediator associations and identify *d* candidate mediators. Second, we randomly split the data into two subsets *B* times and compute the mirror statistic for each candidate mediator within each split. Third, we aggregate the mirror statistics across the *B* splits using inclusion rate aggregation to select mediators under the pre-specified FDR level.

First, we select mediators that are strongly associated with the exposure. Specifically, we obtain the *p*-values in the *X* → *M*_*j*_ (*j* ∈ [*p*]) relationships by fitting linear regression models, and then apply the Benjamini-Hochberg (BH) procedure [21] to select mediators with significant exposure-mediator associations by controlling the FDR at level *q*_1_. After this step, the dimension of candidate mediators is reduced from *p* to *d*. Without loss of generality, we assume this filtering step preserves the first *d* mediators and denote the vector of filtered mediators as 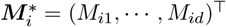.

Second, we randomly split the samples into two groups for *B* times. Denote ***T*** = (*T*_1_, · · ·, *T*_*n*_)^⊤^ as the vector of observed survival outcomes, ***X*** = (*X*_1_, · · ·, *X*_*n*_)^⊤^ as the vector of exposures, 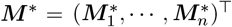 as the design matrix of filtered mediators, and ***Z*** = (***Z***_1_, · · ·, ***Z***_*n*_)^⊤^ as the design matrix of baseline covariates. In the *b*th data split, we randomly split the *n* samples into two groups, Group I and Group II, denoted as (***T*** ^1,*b*^, ***X***^1,*b*^, ***M*** ^∗1,*b*^, ***Z***^1,*b*^) and (***T*** ^2,*b*^, ***X***^2,*b*^, ***M*** ^∗2,*b*^, ***Z***^2,*b*^). In Group I, we fit a Cox proportional hazards model with the MCP and obtain the coefficients 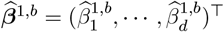, and we denote the set of mediators with non-zero coefficients as 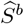. In Group II, we fit a Cox proportional hazards model using only mediators in 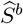 to obtain the corresponding coefficients 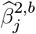; for mediator 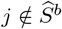, we set 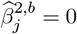. Therefore, we can obtain the vector of coefficients 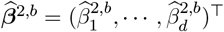 in Group II. We then combine the two vectors of coefficients 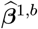 and 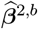 by constructing the mirror statistics 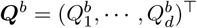 as

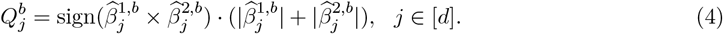

Third, we select mediators by aggregating the mirror statistics over *B* data splits. According to Dai et al. [28], if the *j*th mediator-outcome association *β*_*j*_ = 0, then the mirror statistic 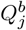 is symmetric about 0; otherwise, 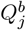 is likely to be a positive and relatively large value. Therefore, if a mediator has large and positive mirror statistics across *B* data splits, then this mediator is more likely to have a non-zero mediator-outcome association. According to Meinshausen et al. [33] and Dai et al. [28], the following two procedures can be used to aggregate the mirror statistics over *B* data splits and then select the mediators with non-zero mediator-outcome associations.

The first aggregation procedure calculates an importance score, named inclusion rate [28], for each mediator over *B* data splits, and selects mediators with large inclusion rates as true mediators, thus we refer to this procedure as the “inclusion rate aggregation”. Specifically, in the *b*th data split, if we select mediators whose mirror statistics are no less than a positive threshold *τ >* 0, then an estimate of the false discovery proportion (FDP) is given by

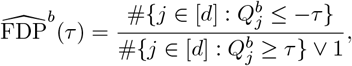

where *a*_1_ ∨*a*_2_ = max(*a*_1_, *a*_2_) for two real numbers *a*_1_ and *a*_2_. For a target FDR control level *q*_2_ ∈ (0, 1), we then choose the data-dependent threshold 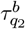 as

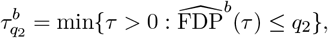

and the selection set of mediators in the *b*th data split is 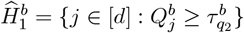. Then we calculate the inclusion rate for each mediator *j* ∈ [*d*] as

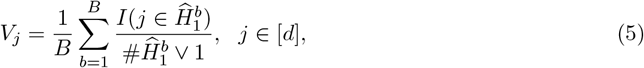

where *I*(·) is the indicator function such that *I*(*A*) = 1 if event *A* happens and *I*(*A*) = 0 if event *A* does not happen. From equation (5), *V*_*j*_ measures the importance of the *j*th mediator across *B* data splits. For example, if the *j*th mediator is selected in each data split, and each data split selects 10 mediators, then *V*_*j*_ = 1*/*10. Therefore, a large value of *V*_*j*_ indicates that the *j*th mediator is more likely to have a non-zero mediator-outcome association. We sort the inclusion rates such that 0 ≤ *V*_(1)_ ≤ *V*_(2)_ ≤ · · · ≤ *V*_(*d*)_, and find the largest index *l* ∈ [*d*] such that *V*_(1)_ + · · · + *V*_(*l*)_ ≤ *q*_2_. Then, the inclusion rate aggregation procedure selects the mediators as

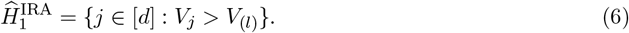

The second aggregation procedure is referred to as the “quantile aggregation”, which is adopted in CoxMKF [27]. Based on the mirror statistic 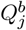 in the *b*th data split, the quantile aggregation calculates the following intermediate statistic 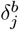 for mediator *j*:

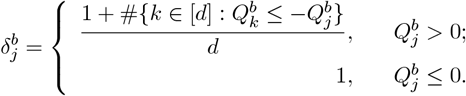

A small value of 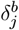 indicates that the *j*th mediator has a strong mediator-outcome association in the *b*th data split. Following Meinshausen et al. [33] and Tian et al. [27], we calculate the Multiple Cox Statistic (MCS) for each mediator:

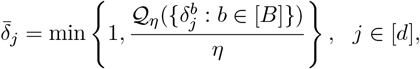

where *η* is the pre-specified quantile point with a recommended value *η* = 0.05 [27], and 𝒬_*η*_(·) denotes the *η*-quantile function. The MCS 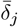 is another importance measurement that is different from the inclusion rate *V*_*j*_. Specifically, a small value of 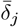 indicates that the *j*th mediator is likely to have a non-zero mediator-outcome association across *B* data splits. We then sort the MCSs such that 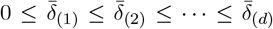, and apply the BH procedure to choose a data-dependent threshold to control the FDR at level *q*_2_:

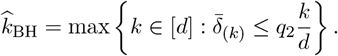

Then, the quantile aggregation procedure selects the mediators as

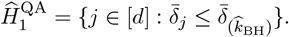

We compare inclusion rate aggregation and quantile aggregation with extensive simulations in Section 3.3. Empirically, we find that the inclusion rate aggregation in CoxMDS maintains finite-sample FDR control even for highly correlated mediators, whereas the quantile aggregation in CoxMDS might have inflated FDR when mediators are highly correlated. Accordingly, we recommend using CoxMDS with inclusion rate aggregation as the default.

## 3 Simulation Studies

In this section, we present extensive simulation studies to assess the finite-sample performance of the proposed CoxMDS framework. In particular, we investigate the following aspects:

1. the impact of the sample-split proportion and the number of splits;
2. the performance of CoxMDS under correlated mediators;
3. the performance of CoxMDS under non-Gaussian mediators.

Each simulation setting is based on 500 replications. In the second and third settings, we compare the performance of CoxMDS with that of CoxMKF. We set *q*_1_ = 0.2 in the screening step and *q*_2_ = 0.1 as the target FDR level. Let 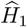 denote the selection set of mediators. We then evaluate CoxMDS and CoxMKF in terms of the false discovery rate (FDR) and the true positive proportion (TPP):

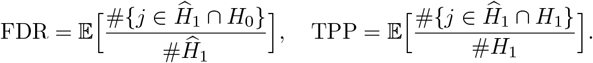

We generate the simulated data as follows. The binary exposure *X*_*i*_ is generated from a Bernoulli distribution with success probability 0.6 as Ber(0.6), and the direct effect of the exposure on the outcome is set to *γ* = 0.5. The covariates are ***Z***_*i*_ = (*Z*_*i*1_, *Z*_*i*2_)^⊤^, where *Z*_*i*1_ is a binary covariate generated from a Bernoulli distribution with success probability 0.3 as Ber(0.3), and *Z*_*i*2_ is a continuous covariate generated from a uniform distribution as U(0, 1). The coefficients of the covariates on the outcome are ***θ*** = (0.3, −0.2)^⊤^. As for the mediators, in the first and second simulation settings, mediators are generated according to model (1), where the coefficients ***ϑ*** = (0.3, −0.2)^⊤^, the error terms follow a multivariate normal distribution with a Toeplitz covariance matrix whose (*i, j*)-th entry is *ρ*^|*i*−*j*|^, and *ρ* ∈ [0, 1) is used to control the correlations among mediators. In the third simulation setting, mediators are generated from beta distributions, as described in Section 3.3. The true event time *D*_*i*_ is generated from the exponential model *λ*_*i*_(*t*|*X*_*i*_, ***Z***_*i*_, ***M***_*i*_) = 0.5 exp {*γX*_*i*_ + ***θ***^⊤^***Z***_*i*_ + ***β***^⊤^***M***_*i*_ }. The censoring time *C*_*i*_ is generated from an exponential distribution with parameter *c*_0_, and the average censoring rate is controlled at 30% or 60% by properly choosing *c*_0_. The observed survival time *T*_*i*_ is generated as *T*_*i*_ = min(*D*_*i*_, *C*_*i*_). For ***α*** and ***β***, we fix the first ten elements of ***α*** to be *κ*_*α*_ · (1, 1, 1, 1, −1, −1, −1, −1, 1, −1), the first twelve elements of ***β*** to be *κ*_*β*_ · (1, 1, 1, 1, 1, 1, 1, 1, 0, 0, 1, −1), where *κ*_*α*_ and *κ*_*β*_ are two scaling factors to prevent the TPP from being too close to 1. The rest elements of ***α*** and ***β*** are set to 0. Therefore, the set of true mediators is *H*_1_ = {1, 2, · · ·, 8}.

### 3.1 Choice of the sample-split proportion and the number of data splits

We evaluate the performance of CoxMDS under a range of sample-split proportions and numbers of data splits. The proportion of samples in Group I ranges from 10% to 90%, and the number of data splits is chosen from {1, 2, 5, 10, 25, 50}. In this simulation setting, we adopt the inclusion rate aggregation and set the target FDR level at *q*_2_ = 0.1. The error terms in model (1) are drawn from a multivariate normal distribution with a Toeplitz covariance matrix, where (*i, j*)-th entry is *ρ*^|*i*−*j*|^ and *ρ* = 0.6. The sample size is set to *n* = 500, and the number of candidate mediators is set to *p* = 2, 000.

As shown in Figure 3, CoxMDS achieves finite-sample FDR control and maintains high TPP with a balanced sample-split proportion and a large number of data splits. For example, with the number of splits *B* = 25 and a 30% censoring rate, a half-half sample-split proportion yields finite-sample FDR control and the highest TPP (0.853) among all sample-split proportions. Moreover, increasing the number of sample splits *B* from 25 to 50 does not yield much power gain. Balancing these trade-offs, we find that a half-half sample-split proportion combined with *B* = 25 data splits achieves finite-sample FDR control, maintains high power, and remains computationally efficient, making it our recommended default for practical applications.

**Figure 3:**
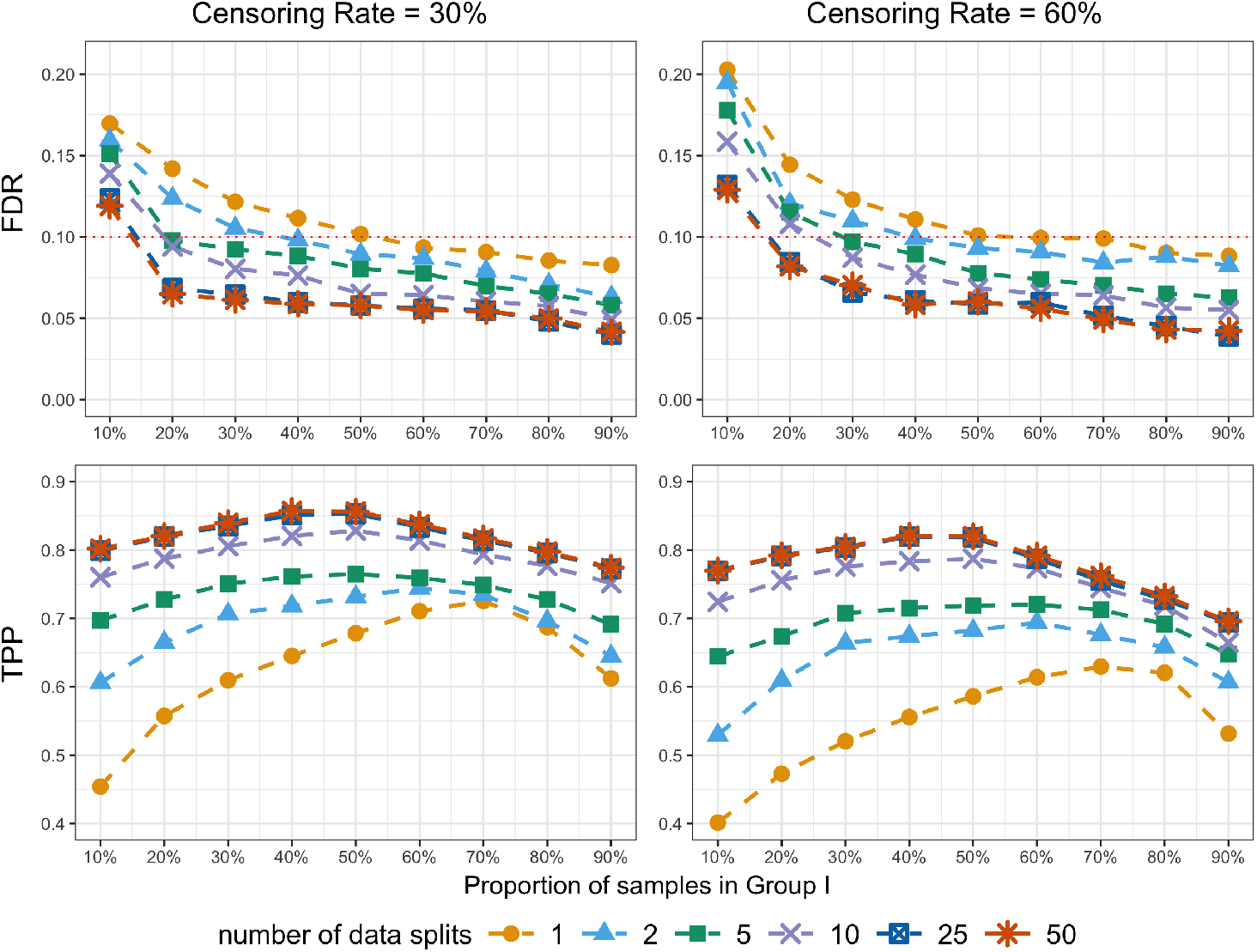
Line plot of the FDR and TPP of CoxMDS against different sample-split proportions and different numbers of data splits. We adopt the inclusion rate aggregation in this simulation. The red dashed line represents the target FDR level *q*_2_ = 0.1.

### 3.2 Comparison of CoxMDS and CoxMKF in correlated mediators

In this simulation, we compare the empirical performance of CoxMDS with CoxMKF under different aggregation procedures (inclusion rate aggregation and quantile aggregation) across different values of correlations among mediators, and the results are presented in Figure 4. Specifically, mediators are generated from model (1), where the error terms are drawn from *N* (0, Σ) with Σ = (Σ_*ij*_)_*p×p*_ = (*ρ*^|*i*−*j*|^)_*p×p*_. We vary the correlation *ρ* ∈ {0, 0.2, 0.4, 0.6, 0.8}, and fix the sample size *n* = 500 and number of candidate mediators *p* = 2, 000. To avoid the power from approaching 1, the scaling factors (*κ*_*α*_, *κ*_*β*_) are set to (0.50, 0.35) in this simulation setting. The default aggregation procedure of CoxMKF is quantile aggregation, but we also implement CoxMKF with inclusion rate aggregation for fair comparison. For quantile aggregation procedure, we adopt the recommended quantile point *η* = 0.05 [27]. We set the target FDR level at *q*_2_ = 0.1 for both CoxMDS and CoxMKF.

**Figure 4:**
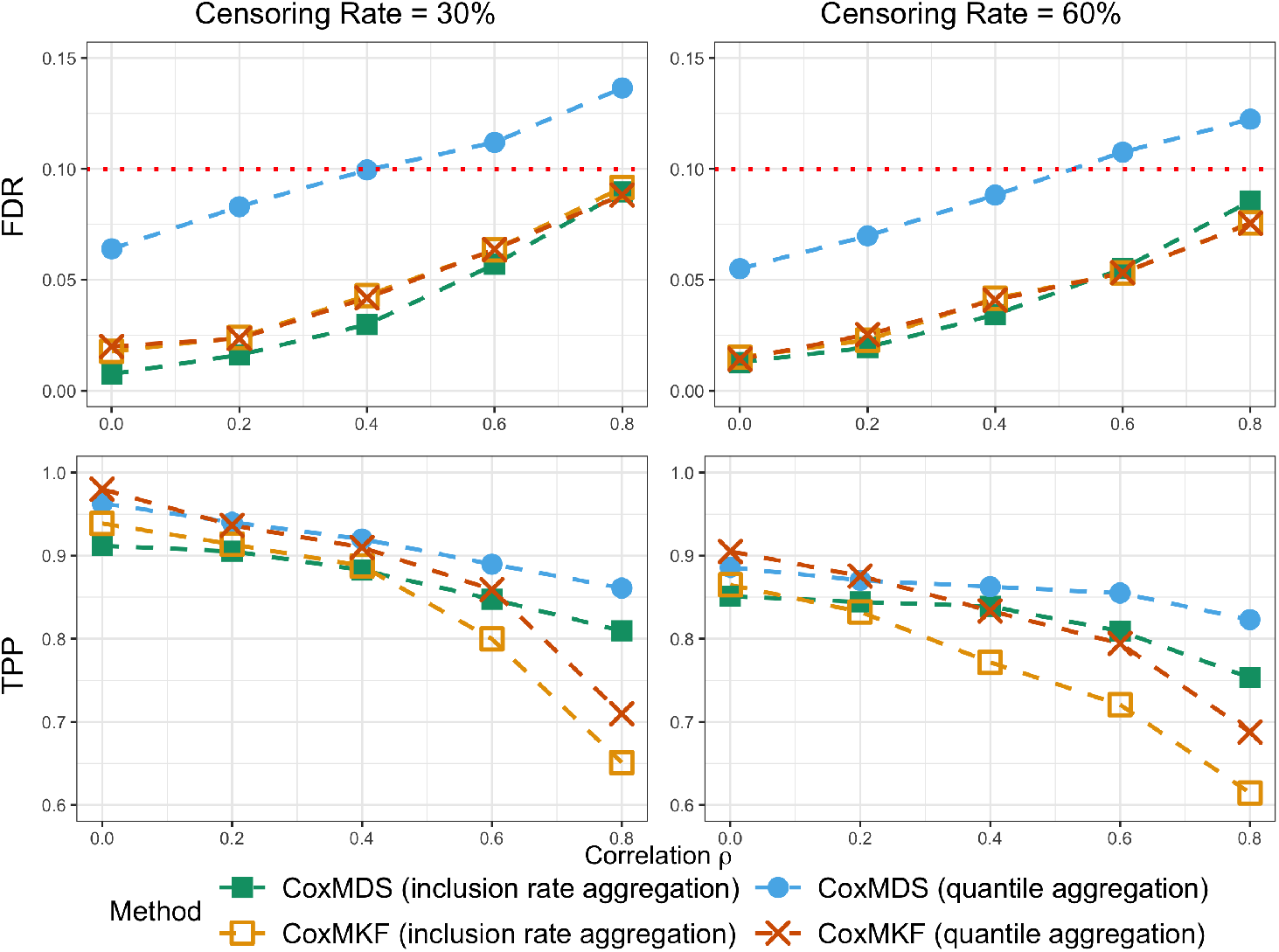
Line plot of the FDR and TPP for CoxMKF and CoxMDS with different values of correlation *ρ* ∈{0, 0.2, 0.4, 0.6, 0.8} and different censoring rates (30% or 60%). The numbers of multiple knockoffs and data splits are both *B* = 25. We adopt a half-half sample-split proportion in CoxMDS. The red dashed line represents the target FDR level *q*_2_ = 0.1.

In this simulation setting, CoxMDS with inclusion rate aggregation not only achieves finite-sample FDR control but also demonstrates greater power than CoxMKF under high mediator correlation. For example, with a censoring rate of 30% and *ρ* = 0.8, the TPP of CoxMDS with inclusion rate aggregation reaches 0.810, compared with 0.651 and 0.710 for CoxMKF under inclusion rate aggregation and quantile aggregation, respectively. The quantile aggregation might result in an inflated FDR when mediators are highly correlated, and thus is not recommended for CoxMDS. These results highlight CoxMDS with inclusion rate aggregation as the preferred method in settings with highly correlated mediators, offering both finite-sample FDR control and higher power. Accordingly, we use inclusion rate aggregation as the default aggregation procedure of CoxMDS in subsequent simulation settings and real data analyses.

### 3.3 Comparison of CoxMDS and CoxMKF in non-Gaussian mediators

In this simulation setting, we investigate the performance of CoxMDS and CoxMKF when the joint distribution of mediators is non-Gaussian. Specifically, we generate the *j*th mediator *M*_*ij*_ from a beta distribution Beta(*a*_*ij*_, *b*_*ij*_), with *a*_*ij*_ = *ϕ*_*j*_ · *µ*_*ij*_ and *b*_*ij*_ = *ϕ*_*j*_ · (1 − *µ*_*ij*_), where *µ*_*ij*_ = 1*/*(1 + exp(−(*c*_*j*_ + *α*_*j*_*X*_*i*_ + ***ϑ***^⊤^***Z***_*i*_)) is the mean of *M*_*ij*_, and *ϕ*_*j*_ *>* 0 is the precision parameter that controls the variance of *M*_*ij*_ and is drawn from an exponential distribution Exp(1). We set the sample size *n* = 500, the number of mediators *p* = 2, 000, the target FDR level *q*_2_ = 0.1, and the scaling factors (*κ*_*α*_, *κ*_*β*_) = (0.90, 0.80). We present the FDR and TPP of CoxMDS and CoxMKF with inclusion rate aggregation procedure in Table 1.

**Table 1:** FDR and TPP for CoxMKF and CoxMDS with different censoring rates (30% or 60%) when mediators are generated from beta distributions. The numbers of multiple knockoffs and data splits are both *B* = 25. The half-half sample-split proportion is adopted in CoxMDS. The target FDR level is set to *q*_2_ = 0.1.

|  | Censoring Rate=30% |  | Censoring Rate=60% |  |
| --- | --- | --- | --- | --- |
|  | FDR | TPP | FDR | TPP |
| CoxMDS | 0.076 | 0.871 | 0.089 | 0.813 |
| CoxMKF | 0.042 | 0.832 | 0.056 | 0.762 |

CoxMDS maintains finite-sample FDR control and achieves higher TPP than CoxMKF in this non-Gaussian setting. For instance, with a 60% censoring rate, the TPP of CoxMDS reaches 0.813, compared with 0.762 for CoxMKF. This performance gap likely arises because CoxMKF relies on knockoffs generated from a multivariate Gaussian distribution, which reduces its power when the candidate mediators are non-Gaussian. In contrast, CoxMDS does not impose distributional assumptions on the mediators, making it more powerful than CoxMKF in non-Gaussian settings.

## 4 Real Data Applications

### 4.1 Identifying CpG Sites Mediating the Effect of Smoking on Lung Cancer Survival

Lung cancer is the most common cancer among men and the third most common among women, with approximately 85% of cases classified as non-small cell lung cancer (NSCLC) [34]. Improving patient survival requires both effective therapies and early diagnosis [10]. Tobacco smoking, the major risk factor for lung cancer, has been shown to induce widespread alterations in DNA methylation [35]. In this section, we apply the proposed CoxMDS method to the TCGA lung cancer dataset [36], which includes patients with lung squamous cell carcinoma and lung adenocarcinoma. Our goal is to identify CpG sites that mediate the effect of smoking on lung cancer survival.

In this dataset, DNA methylation profiles were measured using the Illumina Infinium Human-Methylation450 BeadChip array, and methylation levels were summarized as *β*-values [36]. The exposure is defined as smoking status (smoker vs. non-smoker) at initial diagnosis. The true survival time is the number of days from diagnosis to death, and the censoring time is the number of days from diagnosis to last follow-up. We include age, sex, cancer stage, and radiotherapy status as co-variates. Following Tian et al. [27], we excluded patients with missing survival time, smoking status, or covariates, yielding a final dataset of 754 patients aged 33–90 years with measurements on 365,306 CpG sites. The median survival time is 1,632 days, with a censoring rate of 60%. To explore the distributional properties of the DNA methylation data, we randomly selected four CpG sites and plotted their Pearson correlation heatmap and density curves (Figure 5). These plots reveal that CpG sites are highly correlated and often exhibit non-Gaussian distributions, underscoring the suitability of CoxMDS, which is designed to accommodate both correlated and non-Gaussian mediators.

**Figure 5:**
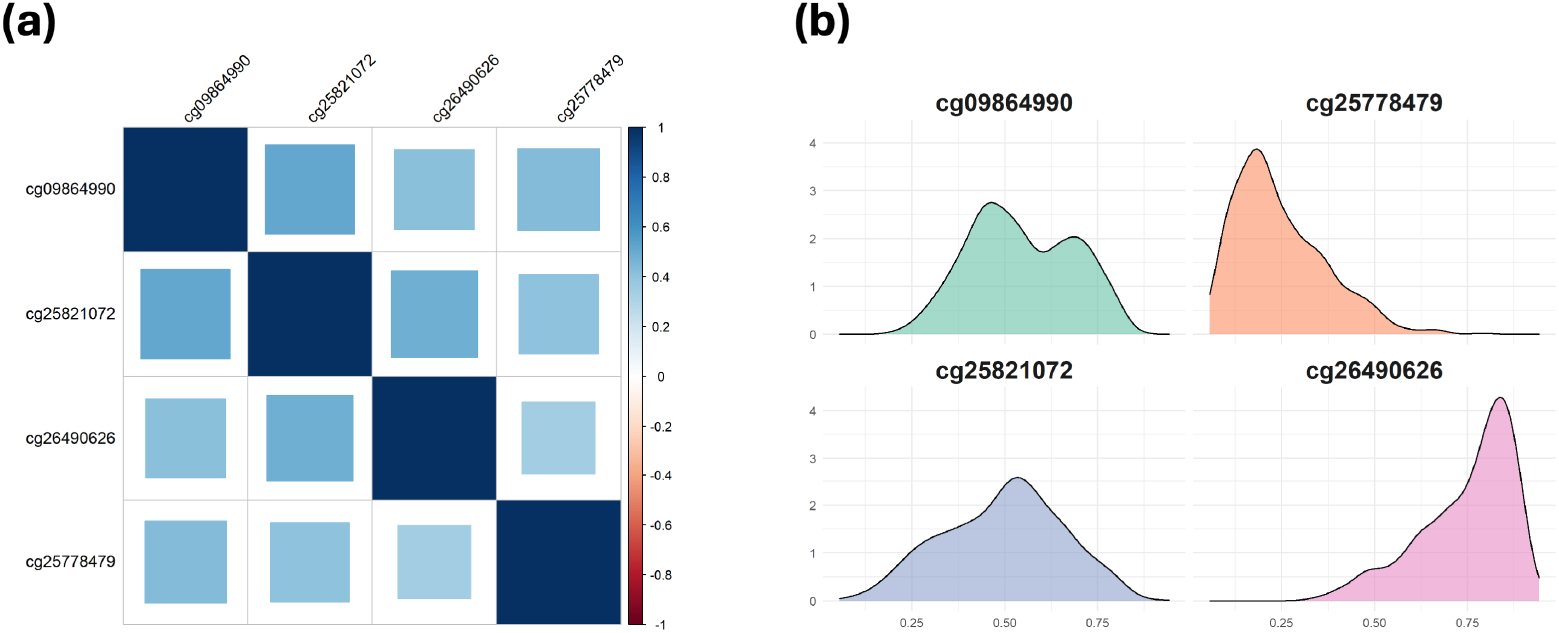
(a) Pearson correlation heatmap and (b) density plots of four DNA methylation CpG sites in the TCGA lung cancer cohort study.

For CoxMDS, we set *q*_1_ = 5 *×* 10^−4^ as the threshold in the BH procedure for candidate mediator filtering, *q*_2_ = 0.1 as the target FDR level, and *B* = 25 as the number of data splits. CoxMDS identifies eight DNA methylation CpG sites, and the effect estimates and corresponding genes are presented in Table 2. All four CpGs previously reported by CoxMKF [27] are also identified by CoxMDS, while four additional CpGs (cg01023787, cg06320150, cg12823072, and cg15953187) are uniquely identified by CoxMDS. This finding is consistent with our simulation results, where CoxMDS demonstrates greater power in detecting correlated and non-Gaussian mediators.

**Table 2:** The identified DNA methylation CpG sites in the TCGA dataset and their estimated effects obtained using CoxMDS with *q*_1_ = 5 *×* 10^−4^, *q*_2_ = 0.1, and *B* = 25. Here, CHR denotes the chromosome.

| CpG | Gene Name | CHR | $\hat{\alpha}$ | $\hat{\beta}$ | $\widehat{\text{NIE}}$ |
| --- | --- | --- | --- | --- | --- |
| cg01023787 | SLC6A18 | 5 | -0.070 | -0.441 | 0.031 |
| cg06320150 | - | 22 | -0.058 | -0.440 | 0.025 |
| cg07690349 | MUC5B | 11 | -0.075 | 1.259 | -0.094 |
| cg12823072 | SCARA5 | 8 | -0.076 | 1.575 | -0.120 |
| cg15953187 | MUC5B | 11 | -0.061 | 0.546 | -0.033 |
| cg21926276 | H19 | 11 | -0.058 | -2.949 | 0.172 |
| cg24129177 | - | 12 | -0.056 | -1.581 | 0.089 |
| cg24200525 | SBF1 | 12 | -0.024 | 5.204 | -0.126 |

Among the newly identified CpGs, cg01023787 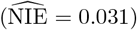 lies in the gene SLC6A18 on chromosome 5. SLC6A18 is located within the 5p15.33 chromosomal region that is frequently gained in early-stage NSCLC, suggesting this locus may participate in early genetic changes associated with the disease [37]. The CpG site cg06320150 also exhibits a positive 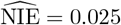 and is associated with the expression of C22orf34, a gene reported to show downregulated expression in lung adenocarcinoma compared with adjacent normal tissue [38]. The CpG site cg12823072 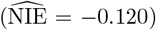 is located in SCARA5 on chromosome 8. SCARA5 acts as a tumor suppressor in NSCLC, with promoter methylation linked to cancer progression [39]. Finally, cg15953187 is located in MUC5B on chromosome 11, the same gene as cg07690349, which is detected by both CoxMDS and CoxMKF. MUC5B is reported as a favorable prognostic biomarker in NSCLC with EGFR mutations and has diagnostic and prognostic significance in lung adenocarcinoma when combined with TTF-1 expression [40]. These DNA methylation CpG sites may provide valuable insights for researchers in designing interventions to improve treatment strategies for lung cancer patients.

### 4.2 Identifying CpG Sites Mediating the Effect of Smoking on Time-to-Conversion to Alzheimer’s Disease

In this section, we apply CoxMDS and CoxMKF to identify CpG sites that might mediate the effect of smoking on time-to-conversion to Alzheimer’s diseases in the Alzheimer’s Disease Neuroimaging Initiative (ADNI) study (https://adni.loni.usc.edu/). The ADNI study investigates Alzheimer’s disease progression through neuroimaging, biomarker measurements, and clinical and cognitive assessments [41]. In this analysis, DNA methylation profiles are measured by the Illumina Infinium HumanMethylationEPIC BeadChip array, and DNA methylation levels are summarized as *M*-values [42]. The exposure is defined as any history of smoking (smoker/non-smoker) during the subject’s life-time. The true survival time is the number of years from recruitment to Alzheimer’s disease diagnosis, and the censoring time is the number of years from recruitment to the last follow-up date. Since the DNA methylation data in the ADNI study are derived from blood samples [42, 43], we estimate the relative proportion of underlying cell types using the R package ENmix [44], and include the estimated cell type proportions as covariates. We also adjust for the following covariates: APOE-*ε*4 allele count (a known genetic risk factor for Alzheimer’s disease [45]), age, years of education, gender, and history of hypertension.

After data preprocessing and quality control [43], we include a total of 611 subjects aged 55–91 years, with 865,859 DNA methylation CpG sites. Of these, 404 subjects are from the ADNI2 phase, while the remaining 207 are from the ADNI GO phase. The censoring rate (i.e., the proportion of subjects without diagnosis of Alzheimer’s disease during the follow-up period) is 66.1%, and the median of survival outcome is 5.10 years. In Figure 6, we present the descriptive statistics of these subjects included in the analysis.

**Figure 6:**
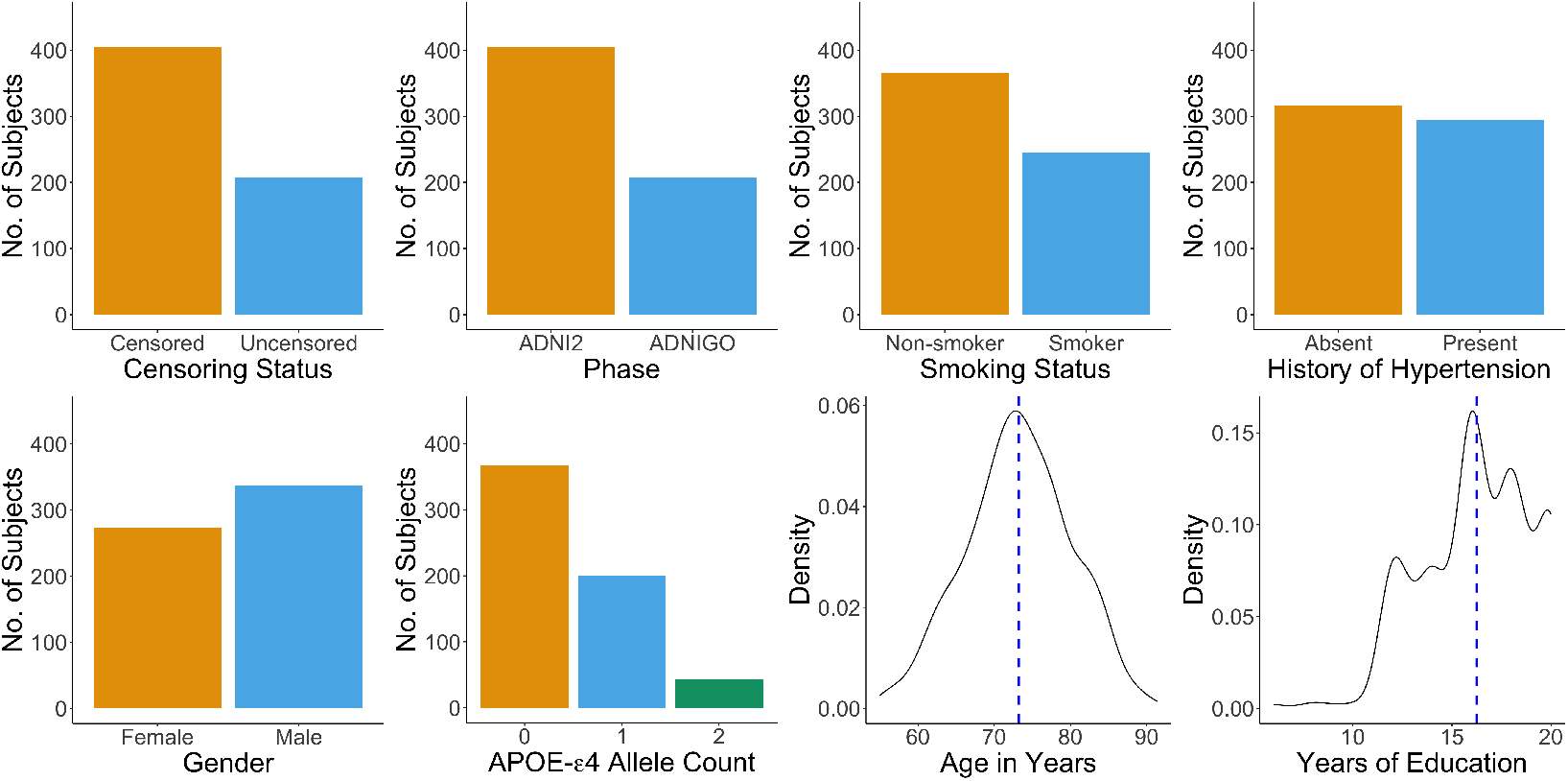
Descriptive statistics of the subjects included in the analysis. Among the 611 subjects, the censoring rate is 66.1%; the proportion of smokers is 40.1%; the proportion of subjects with history of hypertension is 48.1%; the proportion of male subjects is 55.3%; the average APOE-*ε*4 allele count is 0.471; the average age is 73.2 years; and the average years of education is 16.2 years.

For both CoxMDS and CoxMKF, we set *q*_1_ = 0.05 as the threshold for candidate mediator filtering, *q*_2_ = 0.1 as the target FDR level, and *B* = 25 as the number of data splits and multiple knockoffs. CoxMDS identifies two DNA methylation CpG sites (cg06644428 and cg01940273) as mediators, while CoxMKF detects none. This finding aligns with our simulations, which show that CoxMDS achieves higher power than CoxMKF when candidate mediators are highly correlated and exhibit non-Gaussian distributions in finite samples.

We report the two CpG sites identified by CoxMDS in Table 3. Among them, cg06644428 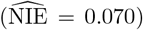 is a known smoking-associated CpG site located in the intergenic region 2q37.1 on chromosome 2 [46]. The CpG site cg01940273 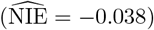, located in gene ALPPL2 on chromosome 2, is also reported as a smoking-associated DNA methylation CpG site [47]. ALPPL2 encodes an alkaline phosphatase protein, and altered plasma levels of alkaline phosphatase are associated with central nervous system injury and Alzheimer’s disease progression [48]. Therefore, these DNA methylation CpG sites may serve as potential targets for future research aimed at understanding disease mechanisms and developing interventions for Alzheimer’s disease.

**Table 3:** The identified DNA methylation CpG sites in the ADNI dataset and their estimated effects using CoxMDS with pre-specified *q*_1_ = 0.05, *q*_2_ = 0.1, and *B* = 25. CHR stands for the chromosome.

| CpG | Gene Name | CHR | $\hat{\alpha}$ | $\hat{\beta}$ | $\widehat{\text{NIE}}$ |
| --- | --- | --- | --- | --- | --- |
| cg06644428 | - | 2 | -0.391 | -0.180 | 0.070 |
| cg01940273 | ALPPL2 | 2 | -0.118 | 0.319 | -0.038 |

## 5 Conclusion

In this paper, we propose CoxMDS, a novel xMediation analysis method in causal genomics, to address the challenge of controlling finite-sample FDR in high-dimensional mediator settings with a survival outcome. By integrating multiple data splitting with high-dimensional mediation analysis, CoxMDS achieves both finite-sample FDR control and high detection power, even when mediators exhibit medium-to-high correlations or follow non-Gaussian distributions. Extensive simulation studies demonstrate that CoxMDS achieves finite-sample FDR control and high detection power even when mediators are highly correlated or non-Gaussian. In real data applications, CoxMDS identifies eight DNA methylation CpG sites that might mediate the effect of smoking on lung cancer survival in the TCGA cohort, and two CpG sites that might mediate the effect of smoking on time-to-conversion to Alzheimer’s disease in the ADNI study.

CoxMDS offers two key advantages over competing methods. First, compared with CoxMKF, CoxMDS maintains finite-sample FDR control while achieving higher power in scenarios with correlated and/or non-Gaussian mediators, making it especially suitable for genome-wide DNA methylation data where co-methylation and non-Gaussian distributions are common. Second, CoxMDS is computationally more efficient than CoxMKF, as it avoids generating knockoff copies of mediators and instead uses random data splitting, which is faster and provides practical scalability for high-dimensional mediation analyses. For example, on the TCGA dataset, the average running times of CoxMDS and CoxMKF are 26.25 minutes and 27.77 minutes, respectively, on a server equipped with an Intel Xeon Silver 4116 CPU and 64 GB RAM. In CoxMDS, most of the computation time is spent in the screening step (25.89 minutes), while the data-splitting step requires only 0.36 minutes, making it more efficient than knockoff generation (1.88 minutes). To facilitate its use, we also provide a user-friendly R package, `CoxMDS`.

CoxMDS opens several avenues for future development. First, we will extend the framework to accommodate accelerated failure time models. Second, we plan to generalize the design from a single exposure–outcome pair to settings with multiple, potentially correlated exposures. Third, incorporating time-varying mediators with repeated measurements will allow longitudinal mediation analyses.

## Key Points

- DNA methylation data often display non-Gaussian distributions and strong correlations, creating challenges for mediator selection and finite-sample false discovery rate (FDR) control.
- We propose CoxMDS, a multiple-data-splitting approach with Cox proportional hazards models for mediator selection in high-dimensional mediation analysis with survival outcomes.
- CoxMDS guarantees finite-sample FDR control and demonstrates higher power than competing methods with highly correlated or non-Gaussian mediators.
- This framework can be broadly applied to other types of omics data, including RNA-seq, proteomics, and metabolomics.

## Code Availability

The CoxMDS method is implemented in an open source R package, which is freely available at https://github.com/MinhaoYaooo/CoxMDS.

## Data Availability

The lung cancer dataset is obtained from The Cancer Genome Atlas (TCGA) program and is publicly available through the UCSC Xena Browser at https://xenabrowser.net/datapages/. The Alzheimer’s disease dataset is obtained from the Alzheimer’s Disease Neuroimaging Initiative (ADNI) and is available upon application at https://adni.loni.usc.edu/.

## Notes

### Competing Interest Statement

The authors have declared no competing interest.

### Summary of Updates

We have revised certain phrases for clarity.

